# Improving GWAS discovery and genomic prediction accuracy in Biobank data

**DOI:** 10.1101/2021.08.12.456099

**Authors:** Etienne J. Orliac, Daniel Trejo Banos, Sven E. Ojavee, Kristi Läll, Reedik Mägi, Peter M. Visscher, Matthew R. Robinson

**Affiliations:** Scientific Computing and Research Support Unit, University of Lausanne, Lausanne, Switzerland; Department of Quantitative Biomedicine, University of Zurich, Zurich, Switzerland; Department of Computational Biology, University of Lausanne, Lausanne, Switzerland; Estonian Genome Centre, Institute of Genomics, University of Tartu, Tartu, Estonia; Institute for Molecular Bioscience, University of Queensland, Brisbane, QLD, Australia; Institute of Science and Technology Austria, Klosterneuburg, Austria

## Abstract

Genetically informed and deep-phenotyped biobanks are an important research resource. The cost of phenotyping far outstrips that of genotyping, and therefore it is imperative that the most powerful, versatile and efficient analysis approaches are used. Here, we apply our recently developed Bayesian grouped mixture of regressions model (GMRM) in the UK and Estonian Biobanks and obtain the highest genomic prediction accuracy reported to date across 21 heritable traits. On average, GMRM accuracies were 15% (SE 7%) greater than prediction models run in the LDAK software with SNP annotation marker groups, 18% (SE 3%) greater than a baseline BayesR model without SNP markers grouped into MAF-LD-annotation categories, and 106% (SE 9%) greater than polygenic risk scores calculated from mixed-linear model association (MLMA) estimates. For height, the prediction accuracy *R*^2^ was 47% in a UK Biobank hold-out sample, which was 76% of the estimated 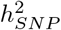. We then extend our GMRM prediction model to provide MLMA SNP marker estimates for GWAS discovery, which increased the independent loci detected to 7,910 in unrelated UK Biobank individuals, as compared to 5,521 from BoltLMM and 5,727 from Regenie, a 43% and 38% increase respectively. The average *χ*^2^ value of the leading markers was 34% (SE 5.11) higher for GMRM as compared to Regenie, and increased by 17% for every 1% increase in prediction accuracy gained over a baseline BayesR model across the traits. Thus, we show that modelling genetic associations accounting for MAF and LD differences among SNP markers, and incorporating prior knowledge of genomic function, is important for both genomic prediction and for discovery in large-scale individual-level biobank-scale studies.

---

As biobank data sets increase in size, it is important to understand the factors limiting the prediction of phenotype from genotype. Alongside others, we have recently shown that genomic prediction accuracy can be improved through the use of random effects models that incorporate prior knowledge of genomic annotations, and which allow for differences in the variance explained by SNP markers depending upon their linkage disequilibrium (LD) and their minor allele frequency (MAF) [1–7]. Improvements in prediction accuracy should also translate into greater GWAS discovery power. Mixed-linear models of association (MLMA) are commonly applied in GWAS studies in a two-step approach, where a random effects model is first used to estimate leave-one-chromosome-out (LOCO) genetic values, and these are then used in a second marginal regression coefficient estimation step. Theory suggests that the test statistics obtained in the MLMA second step depend upon the accuracy of the LOCO genomic predictors produced from the first-step. Current MLMA implementations use either a blocked ridge regression model [8], a REML genomic relationship model [9], or a Bayesian spike-and-slab model [**?**] within the first step.

Here, we improve the computational implementation of our recently developed Bayesian grouped mixture of regressions model (GMRM), which estimates genetic marker effects jointly, but with independent marker inclusion probabilities and independent 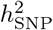 parameters across LD, MAF, and functional annotation groups (see Methods). This allows us to apply the model to 21 traits in the UK Biobank to test for prediction accuracy improvements over existing approaches. We then extend the model to provide MLMA SNP marker association estimates to test whether improved prediction accuracy translates to improved GWAS discovery as compared to existing MLMA approaches.

In an analysis of 428,747 UK Biobank individuals genotyped at 2,174,071 high-quality imputed SNP markers (see Methods), we show that our GMRM model improves 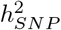 estimation by up to 35.6% (average of 9.09%, SD, 7.9%) as compared to a REML model estimating a single 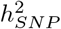 parameter (Figure 1a). The 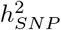 estimate sets the upper bound for prediction accuracy in an independent sample and the higher 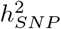 estimates obtained by GMRM likely stem from the fact that genomic annotations differ greatly in the proportion of variance attributable to them, adjusted by that expected given the number of markers included within the model which map to each region (Supplementary Figure 1). In a UK Biobank hold-out sample of 30,000 individuals, a GMRM yields higher prediction accuracy than previously published estimates that we are aware of for all traits (Figure 1b). A GMRM improves over a baseline BayesR model implimented in our software that assumes that markers come from a mixture of four normal distributions and a dirac spike at zero, by and average of 18% (SE 3%, Figure 1c). We determined the best possible prediction accuracy obtained from the LDAK software [4], using either the BLD-LDAK annotations, or the same annotation groups used by the GMRM. We find that LDAK improves prediction accuracy over a BayesR model in many cases (Figure 1c), but that prediction accuracy was generally lower than that obtained by a GMRM (Figure 1b), with a GMRM improving prediction accuracy over the best LDAK model by an average of 15% (SE 7%).

**Figure 1.**
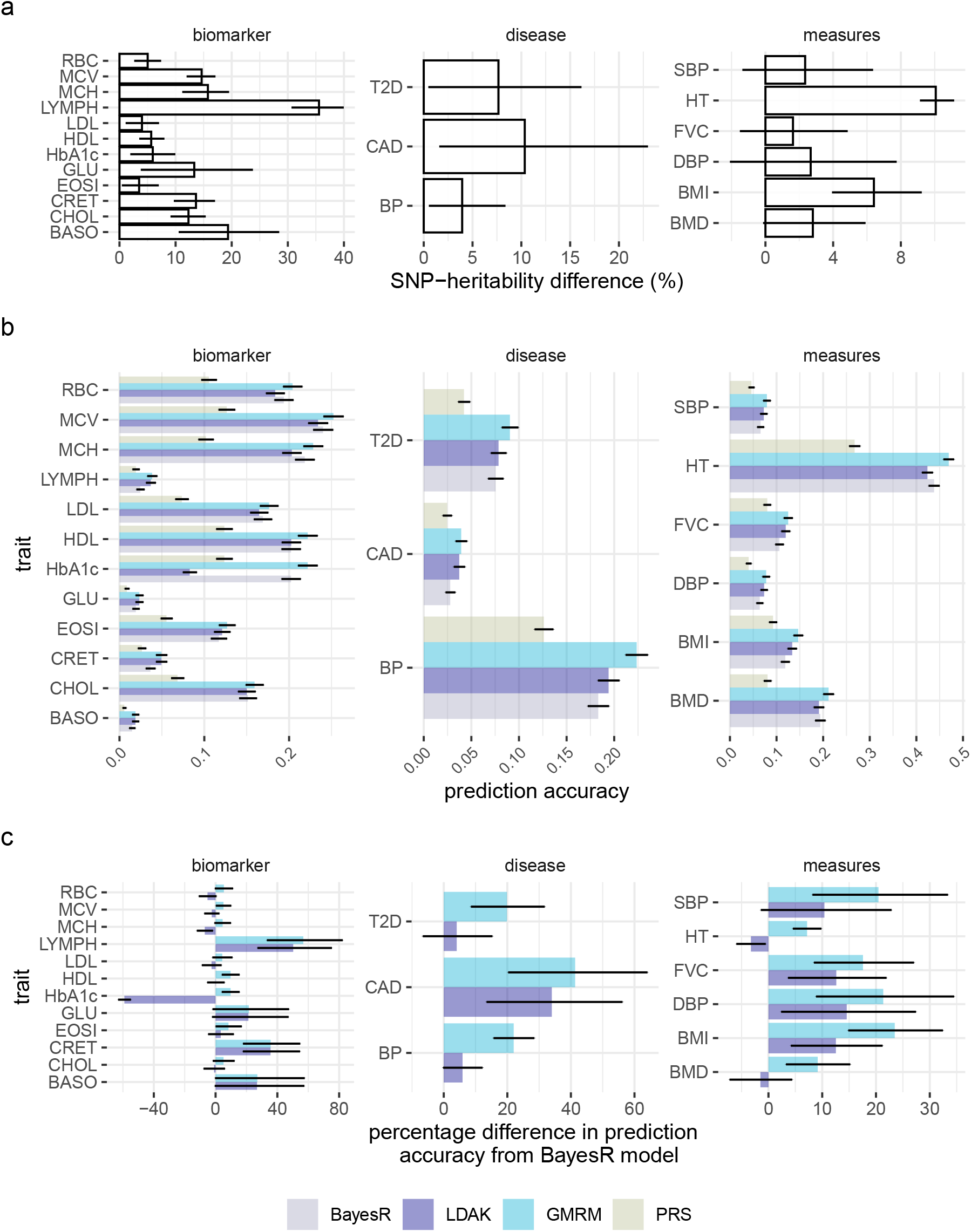
SNP-heritability estimation and prediction accuracy of a GMRM. (a) SNP heritability estimates from a MAF-LD-annotation GMRM as a percentage difference from those obtained from a single component REML analysis across 21 traits. (b) Prediction accuracy obtained by GMRM for the 21 traits as compared to those obtained from the best individual-level LDAK prediction model (LDAK), a BayesR model with five mixture groups (BayesR), or polygenic risk scores calculated using boltLMM SNP marker effects (PRS). (c) Gives the prediction accuracy of LDAK and GMRM models as a percentage difference from the accuracy obtained from the BayesR model. Error bars in give 95% confidence intervals. Full trait code descriptions are given in Supplementary Table 1.

We achieve over 75% of the 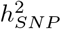 for height, and over 50% for 12 of the traits in the UK Biobank hold-out sample (Figure 2a). The accuracy obtained is higher than that expected from theory by up to 12.5% (mean 4.1%, SD 3.4%, given ridge-regression assumptions and 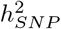 estimates, Figure 2b). We meta-analysed the posterior mean effect sizes obtained from the UK Biobank, with those obtained from a GMRM analysis of 105,000 Estonian Genome Centre participants, and then predicted into an Estonian hold-out sample of 20,000 individuals, improving prediction of body mass index (BMI, prediction accuracy of 16.1%) and cardiovascular disease (CAD, prediction accuracy of 7%) over the accuracy obtained in the UK Biobank hold-out sample (Figure 2c). Previous results have highlighted the lack of transfer of genomic predictors across populations, and here we achieved reduced prediction accuracy for high blood pressure (BP) and type-2 diabetes (T2D) diagnoses in Estonia as compared to the UK Biobank. Thus, while these results highlight the potential of cross-biobank meta-analyses to improve genetic predictors, they also highlight the likely trait dependency of applicability of predictors across different health systems. Nevertheless, we show high stratification across both populations of early-onset risk groups with individuals in the top 1% of predicted genetic values having 7 times (95% CI 4-9) higher risk of cardiovascular disease (CAD), 8 times (95% CI 6-11) higher risk of high blood pressure (BP) in the UK Biobank hold-out sample, and 4 times (95% CI 3-8) higher risk of type-2 diabetes (T2D) prior to 60 years of age, as compared to the rest of the population (Figure 2d). Thus, we show how Bayesian posterior SNP effects size estimates can be meta-analysed across studies to improve identification of individuals at high risk of early common disease onset.

**Figure 2.**
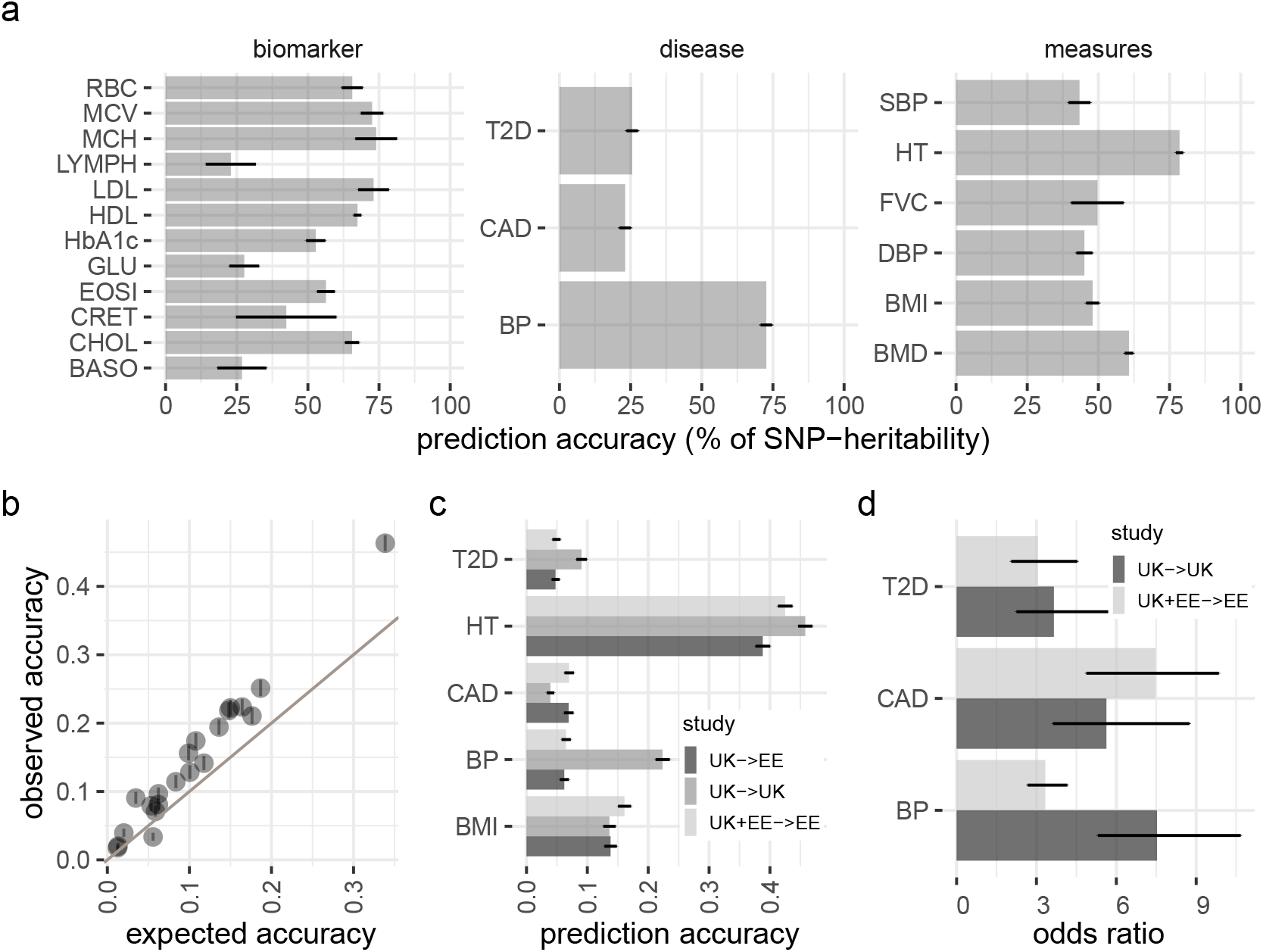
Prediction accuracy of GMRM in UK and Estonian Biobanks. (a) Prediction accuracy of the GMRM effects sizes as a percentage of their upper bound (the SNP-heritability) for 21 traits. (b) Prediction accuracy obtained by GMRM for the 21 traits as compared to that expected from ridge-regression theory. (c) Prediction accuracy obtained using GMRM UK Biobank estimates in UK Biobank hold-out data (UK->UK), GMRM UK Biobank estimates in Estonian data (UK->EE), and UK Biobank and Estonian meta-analysis GMRM estimates in Estonian hold-out data (UK+EE->EE) for five focal traits. (d) Odds ratio for top 1% of the GMRM genetic predictor as compared to all others, within UK->UK and UK+EE->EE for type-2 diabetes (T2D), cardiovascular disease (CAD), and high blood pressure (BP). Error bars in give 95% confidence intervals. Full trait code descriptions are given in Supplementary Table 1.

We modify our GMRM approach to provide leave-one-genomic-region-out, or leave-one-chromosome-out SNP effect estimates (see Methods), and we applied this approach to the UK Biobank data, with first degree relatives removed to minimise the potential for common environment confounding. We validate this approach in simulations (see Methods), showing that our GMRM MLMA approach yields higher power at associated variants, whilst controlling for pervasive population stratification, but not strong common environment effects (Supplementary Figure 2). We find that GMRM MLMA yields greater association testing power than comparable MLMA methods of BoltLMM [**?**], or Regenie [8] for all traits (Figure 3). The number of independent GWAS loci detected at p-value 5×10^−8^ was 7,910 for GMRM MLMA, as compared to 5,521 from BoltLMM and 5,727 from Regenie, a 43% and 38% increase respectively. At regions identified by all approaches at p-value 5×10^−8^, we find that the difference in the *χ*^2^ values obtained by GMRM MLMA as compared to Regenie, scale with the difference in prediction accuracy obtained in independent samples, a relationship expected by theory (see Methods). The average *χ*^2^ value of the leading markers was 34% (SE 5.11) higher for GMRM MLMA as compared to Regenie, and increased by 17% for every 1% increase in prediction accuracy gained over a baseline BayesR model across the traits. A GMRM approach also provides fine-mapping of the associations (WPPA approach, see Methods) and we fine-map 170 associations to single markers with posterior inclusion probability *PIP* ≥ 0.95, 307 associations to SNP sets of 2-5 markers with *PIP* ≥ 0.95, and 497 to groups of 6-20 markers with *PIP* ≥ 0.95 (Supplementary Figure 3). 60% of the total GMRM MLMA associations fine-mapped regions that contained ≥ 100 SNPs in LD. Thus, while we show that modelling genetic associations accounting for MAF and LD differences among SNP markers, and incorporating prior knowledge of genomic function is important for discovery in large-scale individual-level biobank-scale studies, we highlight how LD in the genome creates difficulty for pinpointing the mechanistic basis of the associations.

**Figure 3.**
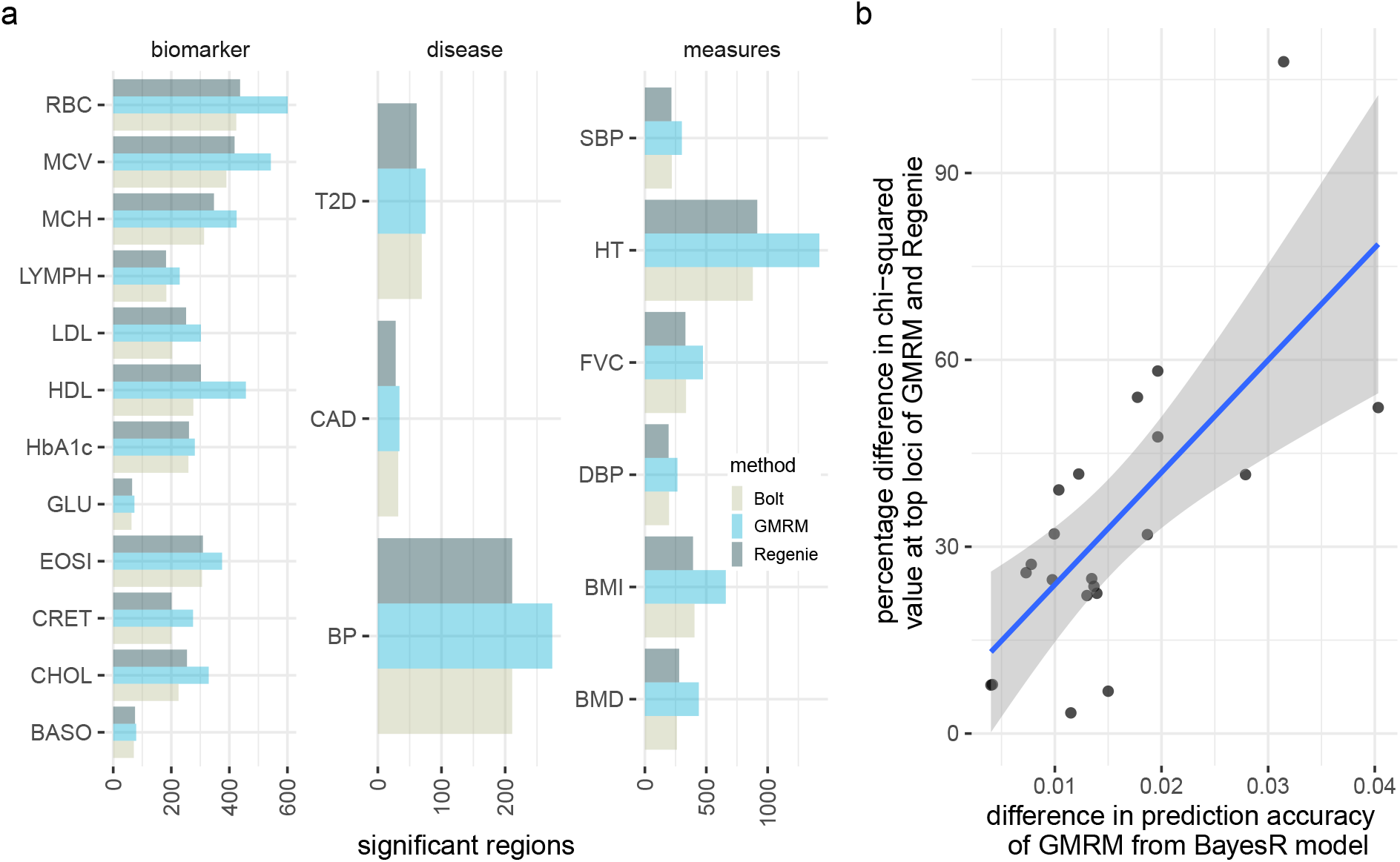
GWAS discovery of GMRM in the UK Biobank. (a) Number of LD independent genomic regions identified at 5×10^−8^ by GMRM, as compared to in boltLMM (bolt) and Regenie (regenie) across 21 traits. (b) For SNP markers identified at 5×10^−8^ by both bolt, regenie, and GMRM, we estimated the percentage difference in *χ*^2^ value between GMRM and regenie and plotted this against the difference in prediction accuracy of GMRM as compared to a BayesR model, to test whether discovery power scales with improve prediction accuracy of using MAF-LD-annotation groups. Shaded area gives the 95% confidence intervals of the regression line. Full trait code descriptions are given in Supplementary Table 1.

There remain important limitations. All of the above analyses were conducted using 2.17 million imputed genetic markers covering all rare high-quality imputed variants (minor allele frequency > 0.001) in a sample of UK Biobank European-ancestry individuals (see Online Methods, Table S1) as a demonstration of the utility of our approach. The portability of polygenic scores across the spectrum of human ethnicity needs to be addressed to avoid polygenic risk stratification that is discriminatory towards groups little represented in currently available genomic data. This is an active research area, and our future work will involve accessing performance of analyses of diverse samples and how transfer learning can improve model estimation across world-wide biobank data. Second, summary statistic approaches have been developed, some of which also account for genomic annotations, MAF and LD when creating genetic predictors [4, 11, 12] and these methods should not be viewed as competing as they are essential for utilising currently available summary data and for situations when individual-level data are not accessible. However, as yet, summary statistics approaches have yet to yield the prediction accuracy obtained in this study [4, 11, 12], and here we have shown how using empirical data from two biobanks can facilitate gains in discovery and polygenic prediction, through a focus on creating powerful and efficient software applications to maximise individual-level data analysis. Thus, we hope that previous consortia analyses can be revisited with a range of improved methodology to facilitate further gains in discovery and polygenic prediction. Our approach is coded in highly optimized C++ code and can accommodate the analysis of multiple traits at once, obtaining trait specific estimates in sub-linear time as the data size increases (see Methods, Supplementary Figures 4 and 5). This work shows that SNP-heritability estimation, association discovery and genomic prediction can be improved simply by better utilizing current data and that as sample size increases, it may become more important to model genetic associations accounting for MAF and LD differences among SNP markers and to incorporate prior knowledge of genomic function.

## Methods

### UK Biobank data

From the measurements, tests, and electronic health record data available in the UK Biobank data [13], we selected 12 blood based biomarkers, 3 of the most common heritable complex diseases, and 6 quantitative measures. The full list of the 21 traits, the UK Biobank coding of the data used, and the covariates adjusted for are given in Supplementary Table 1. For the quantitative measures and blood-based biomarkers we adjusted the values by the covariates, removed any individuals with a phenotype greater or less than 7 SD from the mean (assuming these are measurement errors), and standardized the values to have zero mean and variance 1.

For the common complex diseases, we determined disease status using a combination of information available. For high blood pressure (BP), we used self-report information of whether high blood pressure was diagnosed by a doctor (UK Biobank code 6150-0.0), the age high blood pressure was diagnosed (2966-0.0), and whether the individual reported taking blood pressure medication (6153-0.0, 6177-0.0). For type-2 diabetes (T2D), we used self-report information of whether diabetes was diagnosed by a doctor (2443-0.0), the age diabetes was diagnosed (2976-0.0), and whether the individual reported taking diabetes medication (6153-0.0, 6177-0.0). For cardiovascular disease (CAD), we used self-report information of whether a heart attack was diagnosed by a doctor (3894-0.0), the age angina was diagnosed (3627-0.0), and whether the individual reported heart problem diagnosed by a doctor (6150-0.0) the date of myocardial infarction (42000-0.0). For each disease, we then combined this with primary death ICD10 codes (40001-0.0), causes of operative procedures (41201-0.0), and the main (41202-0.0), secondary (41204-0.0) and inpatient ICD10 codes (41270-0.0). For BP we selected ICD10 codes I10, for T2D we selected ICD10 codes E11 to E14 and excluded from the analysis individuals with E10 (type-1 diabetes), and for CAD we selected ICD10 code I20-I29. Thus, for the purposes of this analysis, we define these diseases broadly simply to maximise the number of cases available for analysis. For each disease, individuals with neither a self-report indication or a relevant ICD10 diagnosis, were then assigned a zero value as a control.

We restricted our discovery analysis of the UK Biobank to a sample of European-ancestry individuals. To infer ancestry, we used both self-reported ethnic background (21000-0) selecting coding 1 and genetic ethnicity (22006-0) selecting coding 1. We also took the 488,377 genotyped participants and projected them onto the first two genotypic principal components (PC) calculated from 2,504 individuals of the 1,000 Genomes project with known ancestries. Using the obtained PC loadings, we then assigned each participant to the closest population in the 1000 Genomes data: European, African, East-Asian, South-Asian or Admixed, selecting individuals with PC1 projection < absolute value 4 and PC 2 projection < absolute value 3. Samples were excluded if in the UK Biobank quality control procedures they (i) were identified as extreme heterozygosity or missing genotype outliers; (ii) had a genetically inferred gender that did not match the self-reported gender; (iii) were identified to have putative sex chromosome aneuploidy; (iv) were excluded from kinship inference; (v) had withdrawn their consent for their data to be used. We used the imputed autosomal genotype data of the UK Biobank provided as part of the data release. We used the genotype probabilities to hard-call the genotypes for variants with an imputation quality score above 0.3. The hard-call-threshold was 0.1, setting the genotypes with probability ≤ 0.9 as missing. From the good quality markers (with missingness less than 5% and p-value for Hardy-Weinberg test larger than 10-6, as determined in the set of unrelated Europeans) we selected those with minor allele frequency (MAF) > 0.0002 and rs identifier, in the set of European-ancestry participants, providing a data set 9,144,511 SNPs. From this we took the overlap with the Estonian Genome centre data described below to give a final set of 8,430,446 markers. For computational convenience we then removed markers in very high LD selecting one marker from any set of markers with LD *R*^2^ ≥ 0.8 within a 1MB window. These filters resulted in a data set with 458,747 individuals and 2,174,071 markers.

We split the sample into training and testing sets for each phenotype, selecting 30,000 individuals that were unrelated (SNP marker relatedness <0.05) to the training individuals to use as a testing set. This provides an independent sample of data with which to access prediction accuracy. For the complex diseases, we randomly select 1000 cases and match to 29,000 controls, again ensuring that these individuals were unrelated to those in the training sample.

### Estonian Biobank data

The Estonian Biobank (EstBB) is a population-based cohort encompassing 20% of Estonia’s adult population (200’000 individuals; 66% females; https://genomics.ut.ee/en/research/estonian-biobank). Individuals underwent microarray-based genotyping at the Core Genotyping Lab of the Institute of Genomics, University of Tartu. 136,421 individuals were genotyped on Illumina Global Screening (GSA) arrays and we imputed the data set to an Estonian reference, created from the whole genome sequence data of 2,244 participants [14]. From 11,130,313 markers with imputation quality score > 0.3, we selected SNPs that overlapped with those selected in the UK Biobank, resulting in a set of 2,174,071 markers.

General data, including basic body measurements, were collected at recruitment. Project-based questionnaires were sent later and filled on a voluntary basis. Health records are regularly updated through linkage with the national Health Insurance Fond and other relevant databases, providing sporadic access to blood biomarker measurements and medical diagnoses [15]. For the genotyped individuals, we had data available for height and body mass index and we removed individuals plus or minus 7SD from the mean and adjusted both phenotypes by the age at enrollment, sex, and the first 20 PCs of the SNP marker data. Prevalent cases of BP, CAD and T2D in the Estonian Biobank cohort were first identified on the basis of the baseline data collected at recruitment, where the information on prevalent diseases was either retrieved from medical records or self-reported by the participant. The cohort was subsequently linked to the Estonian Health Insurance database that provided additional information on prevalent cases (diagnoses confirmed before the date of recruitment) as well as on incident cases during the follow-up. For BP we selected ICD10 code I10, for CAD we selected codes of I20-I29, and for T2D we selected codes E11-E14 and excluded E10. We also split the sample into training and testing sets for each phenotype, selecting 20,000 individuals that were unrelated (SNP marker relatedness <0.05) to the training individuals to use as a testing set. This provides an independent sample of data with which to access prediction accuracy. For the complex diseases, we randomly select 1000 cases and match to 19,000 controls, again ensuring that these individuals were unrelated to those in the training sample.

### Multiple outcome Bayesian grouped mixture of regressions model (GMRM)

We extend the software implementation of our recently developed Bayesian grouped mixture of regression model [1]. In brief, our GMRM model is as follows

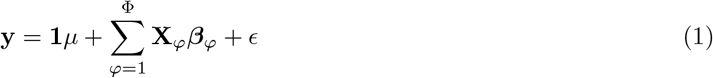

where there is a single intercept term **1***μ* and a single error term *ϵ* but SNPs are allocated into groups (*φ*_1_, ..., *φ*_Φ_), each of which has it’s own set of model parameters 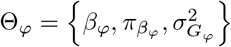. As such, each 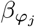 is distributed according to:

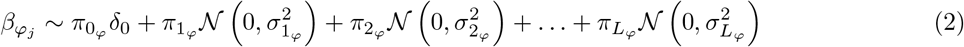

where for each SNP marker group 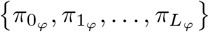 are the mixture proportions and 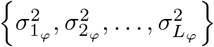 are the mixture-specific variances proportional to

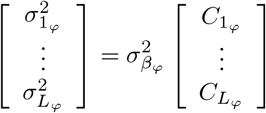

Thus the mixture proportions, variance explained by the SNP markers, and mixture constants are all unique and independent across SNP marker groups. Following our previous work [1], we partition SNP markers into 7 location annotations using the knownGene table from the UCSC browser data, preferentially assigned SNPs to coding (exonic) regions first, then in the remaining SNPs we preferentially assigned them to intronic regions, then to 1kb upstream regions, then to 1-10kb regions, then to 10-500kb regions, then to 500-1Mb regions. Remaining SNPs were grouped in a category labelled “others” and also included in the model so that variance is partitioned relative to these also. Thus, we assigned SNPs to their closest upstream region, for example if a SNP is 1kb upstream of gene X, but also 10-500kb upstream of gene Y and 5kb downstream for gene Z, then it was assigned to be a 1kb region SNP. This means that SNPs 10-500kb and 500kb-1Mb upstream are distal from any known nearby genes. We further partition upstream regions to experimentally validated promoters, transcription factor binding sites (tfbs) and enhancers (enh) using the HACER, snp2tfbs databases (see Code Availability). All SNP markers assigned to 1kb regions map to promoters; 1-10kb SNPs, 10-500kb SNPs, 500kb-1Mb SNPs are split into enh, tfbs and others (un-mapped SNPs) extending the model to 13 annotation groups. Within each of these annotations, we have three minor allele frequency groups (MAF<0.01, 0.01>MAF>0.05, and MAF>0.05), and then each MAF group is further split into 2 based on median LD score. This gives 78 non-overlapping groups for which our model jointly estimates the phenotypic variation attributable to, and the SNP marker effects within, each group. For each of the 78 groups, SNPs were modelled using five mixture groups with variance equal to the phenotypic variance attributable to the group multiplied by constants (mixture 0 = 0, mixture 1 = 0.00001, mixture 2 = 0.0001, 3 = 0.001, 4 = 0.01, 5 = 0.1). The probabilities that markers enter each of the five mixtures (the mixture components) and the variance attributable to the groups are all estimated independently, linked only by a residual updating scheme and thus a regression problem where the number of covariates greatly exceeds the number of measured individuals is broken down into a series of interdependent regressions where the number of covariates within a group is always far less than the total sample size.

We first extend our prediction software to accommodate the analysis of multiple traits simultaneously. While this approach does not yet utilise estimates of the genetic or residual covariance among different outcomes when estimating the SNP effects, a number of coding developments were made to improve speed of the baseline calculations and facilitate the random number generation, vectorisation of the effect size estimation and missing data handling for multiple outcomes. We bench-marked the timing of our software increasing the sample size, marker number, number of traits analysed, and the number of MPI processes, to demonstrate the scalability of our approach as the dimensionality of the data increases. We present these results in Supplementary Figure 4.

We apply the model to the 21 UK Biobank traits described above and use the posterior mean SNP marker effects to predict into the hold-out sample. We repeat the analysis without the MAF-LD-annotation groups, fitting five mixture components, which is equivalent to a BayesR model. We then repeat the analysis again using the individual-level prediction models implemented in LDAK and described in a recent paper [4] with both the BLD-LDAK annotations, and the same annotations used for the GMRM model as described above. We then present the highest prediction accuracy obtained as measured by the correlation between the predictor and the phenotype within the hold-out UK Biobank sample. Finally we also repeated the analysis using boltLMM, selecting an LD pruned (*R*^2^ ≤ 0.01 within 1MB window) subset of markers for the random effect terms and used the marker effects to create a predictor. We present these comparisons within Figure 1. We then predict into the Estonian Biobank data, run the GMRM in the Estonian data, and predict into the Estonian hold-out sample using both UK Biobank model estimates and UK and Estonian Biobank combined estimates. We present these comparisons in Figure 2.

Second, we then extend the software to return the typical fixed effect SNP regression coefficients estimated by other mixed linear model approaches (MLMA). In the first step of running GMRM, the SNP marker effects that we obtain are jointly estimated and thus within each iteration estimation is made accounting for the effects of other markers in both short- and long-range LD. From these estimates, we can obtain partitioned predictors for each focal block *k* of the genome 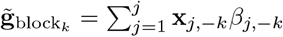, where **x**_*j,−k*_ are the values of SNP markers that are not part of block *k* and *β*_*j,−k*_ their jointly estimated posterior mean effects. These can then be used when testing for association in a second step to yield standard frequentist mixed-model summary statistics, following other approaches. When testing for association of the phenotype with a marker **x**_*j*_ from focal block *k* we consider a simple linear model:

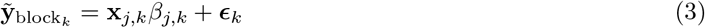

where 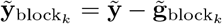 gives the phenotypic residuals where the polygenic effects estimated across the genome other than the focal testing block are adjusted for, **x**_*j*_ is the *j*^*th*^ marker in the focal block, *β*_*j*_ is the OLS estimate for the *j*^*th*^ marker in block *k* and ***ϵ***_*k*_ is the residual error, with 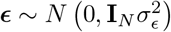 with *M*_*k*_ the number of markers within block *k*.

A t-test statistic is then straight forward to obtain as:

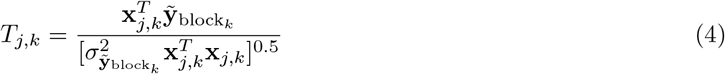

where 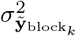 is calculated as 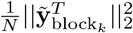 where *N* is the number of individuals. A normal approximation of 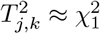 is used to give the p-value. A step-by-step algorithm for this GMRM MLMA approach is given in Algorithm 1.

The test statistic values obtained are an approximation of the mixed-model *χ*^2^ statistic if one were to model the SNP as a fixed effect with a full mixed model equation. This approach follows recent studies (Eq. 23 and 24 of Supplementary Online Material of ref [**?**] and ref [8]), in particular as the *χ*^2^ statistic obtained from boltLMM is equivalent to computing the squared correlations between SNPs being tested and a best linear unbiased predictor, which is the approach taken here. The power of mixed-model association is driven by the fact that focal test SNPs are tested against a ‘denoised’ residual phenotypes, from which other SNP effects estimated by the mixed model have been conditioned out. Here, we expect the SNP marker effect estimation to be improved as the genetic predictor used to residualise the phenotype should have a higher prediction accuracy within independent sample if the true underlying SNP marker effects differ across MAF-LD-annotation groups. While further work is required to optimise this approach for rare diseases using saddle point approximations and to account for the covariance of multiple outcomes, here we wished to simply demonstrate that modelling SNP effects in MAF-LD-annotation groups in the first step yields improved MLMA fixed effect marker estimates in the second step. We verify this approach in a simulation study as described below.

Guided by the simulation study results, we subset the UK Biobank data to individuals that were related at less than first degree relatives (N = 414,055). We then obtained MLMA estimates using GMRM MLMA and we also verified these by placing the LOCO predictors that we obtain from GMRM into the second step of the Regenie software. This enabled a direct comparison with Regenie and we present these results for GMRM MLMA comparing them to those obtained from Regenie on the same data. Furthermore, we also compared to the results obtained by BoltLMM. We present these MLMA results in Figure 3.

### Simulation study

We follow a similar simulation study design as presented in a number of recent studies [9]. From our quality controlled UK Biobank data, we first randomly selected 100,000 individuals with relatedness estimated from SNP markers of ≤ 0.05 and used marker data from chromosomes 1 through 10 with MAF ≥ 0.001 to give 1.36 million SNPs. For the odd chromosomes, we randomly selected 10,000 LD independent SNP markers as causal variants. We simulated effect sizes for the causal variants, **b**, by drawing from a normal distribution with zero mean and variance 0.5/10000. We then scaled the 10,000 causal variant SNP markers to have mean 0 and variance 1 and multiplied them by the simulated marker effect sizes to give genetic values with mean 0 variance 0.5. In previous work, we have extensively explored the ability of our GMRM approach to recover SNP effects sizes and accurately estimate SNP-heritability parameters independently of the relationship of effect sizes, MAF and LD, and so here we simply assume a simple underlying genetic model of randomly selected causal variants and no relationship between effect size and MAF or LD. We then simulate population stratification effects by scaling the loadings of the 100,000 individuals on the first principal component calculated from the genetic data and multiplying the values by 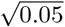, this gives a vector of values, *ps* with variance 0.05. Finally, individual environment values are drawing from a normal distribution with zero mean and variance 1 − var(**Xb**) − var(*ps*), so that the sum of the genetic, *ps*, and environmental values give a phenotype with zero mean and variance 1. We replicate this simulation 10 times, referring to it in the main text and the figures as “random unrelated” causal variant allocation.

Using the same randomly selected individuals, we repeat the simulation but we separate the genetic markers into 4 groups randomly selecting 1000 LD independent exonic markers, 1000 LD independent intronic markers, 1000 LD independent transcription factor binding site markers, and 7000 LD independent markers from other annotation groups. We then sample the marker effects for each of the four groups, drawing from independent normal distributions with zero mean and variance 0.1/1000, 0.25/1000, 0.1/1000, and 0.05/7000 respectively. This gives three annotation groups with larger effect size variance but the same number of causal variants and a total variance explained by the SNP markers var(**Xb**) = 0.5 as the previous simulation setting. We then simulated population stratification effects in the same way and individual environment values by drawing from a normal distribution with zero mean and variance 1 − var(**Xb**) − var(*ps*), so that the sum of the genetic, *ps*, and environmental values give a phenotype with zero mean and variance 1. We also replicate this simulation setting 10 times, referring to it in the main text and the figures as “enrichment unrelated” causal variant allocation.

We then repeat the “random” and the “enrichment” simulation settings, but we change the selection of the individuals used for the simulation. We randomly select 10,000 unique sibling pairs from the UK Biobank data and combine this with 80,000 randomly selected unrelated individuals to give a mixture of relatedness similar to the proportions of related and unrelated individuals in the UK Biobank data. We simulate the same genetic and *ps* values, but we also add a common environmental variance by drawing a value for each sibling pair from a normal distribution with mean 0 and variance 1 and allocated a value of 0 to each of the unrelated individuals. We scale these values by 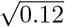 to give a vector of among-family differences, *pe*, with variance 0.12. Finally, individual environment values are drawing from a normal distribution with zero mean and variance 1 − var(**Xb**) − var(*ps*) − var(*pe*), so that the sum of the genetic, *ps*, *pe*, and environmental values give a phenotype with zero mean and variance 1. We replicate this simulation 10 times, referring to it in the main text and the figures as “random related” and “enrichment related” causal variant allocation.

We then analyse this data using bolt-LMM [**?**], fastGWA [9], and GMRM-MLMA and calculate the average *χ*^2^ values (using an approximation that *T*^2^ ≈ *χ*^2^) for the causal variants, which gives a comparative measure of the power of each approach. We also calculate the average *χ*^2^ values (again using an approximation that *T*^2^ ≈ *χ*^2^) obtained from markers on the even chromosomes that contain no causal variants, comparing to the null expectation of 1. These results are presented in Supplementary Figure 2. We find improved power of GMRM-MLMA over other approaches in all settings, with control of the pervasive population stratification. We find only moderate *χ*^2^ inflation when relatives share strong common environment effects that is the same as that obtained by Bolt-LMM. This inflation never increased the false discovery rate (FDR) above 2% in any of the simulations, but was best controlled by a fastGWA approach.

### Utilizing the posterior distribution obtained

We apply our model to each UK Biobank and Estonian Genome Centre data trait, running two short chains for 5000 iterations and combining the last 2000 posterior samples together. We show in Supplementary Figure 5 that the prediction accuracy obtained from our model and the hyperparameter estimates of 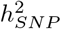 converge within the first 2000 iterations. While obtaining a full posterior distribution with many hundreds of independent samples would require running longer chains, we show that the posterior mean effect size for each SNP that we use for prediction (and thus also in the estimation of the MLMA effect sizes) is well approximated within this run time (Supplementary Figure 5), which is sufficient in this work to assess the prediction accuracy of our approach and to estimate the variance attributable to different genomic regions.

We estimate the proportion of total 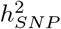 attributable to each genomic annotation and we divide this proportion, by the proportion of SNP markers in the model for this annotation given the total number of SNP markers in the model. This gives an estimate of the enrichment of the marker effects, whereby if the average effect sizes of markers within a given annotation are larger than expected given the number of markers entering the model for that annotation then the value obtained should be greater than 1. Conversely, smaller than expected marker effects will yield values less than 1. We present these results in Figure S1, where we find substantial enrichment of SNP-heritability in intronic regions across traits, and evidence that enrichment differs across traits, with exonic enrichment for height, forced vital capacity, mean corpuscular heamoglobin, low and high density lipoprotein, and blood cholesterol levels (Supplementary Figure 1). Generally, blood based biomarkers show enrichment at proximal promotors, transcription factor binding sites and enhancers, with variation in complex diseases and quantitative traits attributable to distal transcription factor binding sites and enhancers in proportion to that expected given the number of markers in the model (Supplementary Figure 1). SNP markers located greater than 500kb from a gene, explained a far smaller proportion of variance explained then expected given the number of markers which map to the region (Supplementary Figure 1).

We previously presented a window posterior probability of association approach (WPPA) [1]. The WPPA, is estimated by counting the proportion of MCMC samples in which the regression coefficent *β*_*j*_ is greater than a given threshold for at least one SNP *j* in a given genomic window, which can be used as a proxy for the posterior probability that the genomic region contains a causal variant. Because WPPA for a given window is a partial association conditional on all other SNPs in the model, including those flanking the region, the influence of flanking markers on the WPPA signal for any given window will be inversely related to the distance *k* of the flanking markers. Thus, as the number of markers between a causal variant and the focal window increases, the influence of the causal variant on the WPPA signal will decrease and so WPPA computed for a given window can be used to locate associations for that given window, whilst also controlling the false discovery rate. Thus, it represents an approach to fine-mapping association results to groups of SNP markers. Here, we group markers by LD into 341,380 LD independent groups using plink’s clumping procedure, which selects groups of markers (from single SNPs to groups of 100 or more) with LD *R*^2^ ≥ 0.1 within a 1Mb window. For each of these 341,380 SNP groups we calculate the WPPA, defined here as the posterior probability that a group explains at least 0.0001% of the phenotypic variance and we present the number of groups with WPPA ≥ 0.95 in Supplementary Figure 3.

### Comparisons to theory

We note the important distinction between prediction SNP effect sizes that are obtained from mixed-models/penalized regression models, in which a single model is fit where all SNPs are included as random effects, and those obtained from an MLMA model where individual SNP association estimates are typically sought. Here, we sought to make this distinction clear by comparing the prediction accuracy obtained by GMRM and it’s baseline BayesR model, to that obtained from using other random effect models implemented in LDAK, and to predictors created from MLMA estimates to demonstrate that it is inappropriate to create polygenic risk scores from MLMA estimates as compared to random effect models. The expected prediction accuracy in an independent sample under ridge regression assumptions is given by ref [**?**] as:

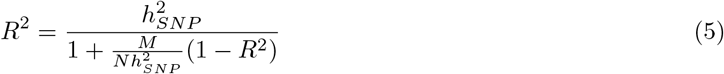

and we use this equation, with the number of markers in the model as a proxy for *M* and the estimates of 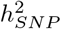 obtained from GMRM, to compare to the prediction accuracy we obtain.

For the common complex disease traits, we place the estimates of the proportion of variance explained by the SNP markers on the liability scale to facilitate comparison with the quantitative measures. The linear transformation of heritability from the observed 0-1 scale, 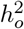, to that of liability scale 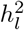 is:

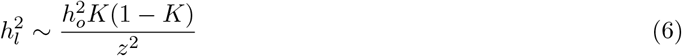

with *K* the population lifetime prevalence and *z* the height of the normal curve at the truncation point pertaining to *K* [17]. We did not observe the population lifetime prevalence of any disease within either population and so we make the assumption that the prevalence in the UK Biobank and Estonian Biobank samples provides a very distant approximation. We scale the estimates of the prediction accuracy to also place these on the liability scale, 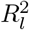, as:

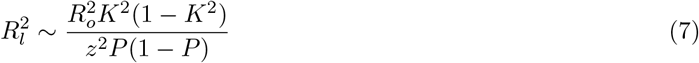

with *P* the proportion of cases in the testing sample. Note that this expression is also an approximation and can result in bias when ascertainment is extreme and heritability on the liability scale is high, though this is expected to be small in practice and negligible for the three disease trait considered here [18].

Previous work [19] has shown that the expectation of the mean *χ*^2^ value at causal SNPs is given by:

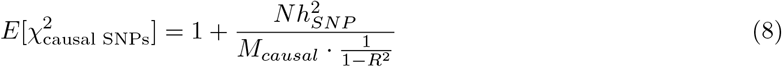

with N the sample size. Thus, under the assumptions of no confounding, no case-control ascertainment-induced confounding, and no pervasive familial relatedness, increased prediction accuracy should yield increased power (*χ*^2^ statistics) at SNPs that are associated with underlying causal variants.

## Author Contributions

MRR and PMV conceived and designed the study. MRR conducted the study with input from EJO, SEO, DTB, KL and RM, with RM providing Estonian Biobank study oversight. EJO and MRR developed the software. MRR wrote the paper. All authors approved the final manuscript prior to submission.

## Acknowledgements

This project was funded by an SNSF Eccellenza Grant to MRR (PCEGP3-181181), and by core funding from the Institute of Science and Technology Austria. PMV acknowledges funding from the Australian National Health and Medical Research Council (1113400) and the Australian Research Council (FL180100072). K.L. and R.M. were supported by the Estonian Research Council grant PRG687. Estonian Biobank computations were performed in the High Performance Computing Centre, University of Tartu.

## Data availability

UK Biobank has approval from the North West Multi-centre Research Ethics Committee (MREC) to obtain and disseminate data and samples from the participants (http://www.ukbiobank.ac.uk/ethics/), and these ethical regulations cover the work in this study. Written informed consent was obtained from all participants. Data from this project were held under UK Biobank project ID 35520. The individual-level genotype and phenotype data are available through formal application to the UK Biobank (http://www.ukbiobank.ac.uk).

The Estonian Biobank data are available upon request from the cohort author RM according to data access protocols to researchers with approval from Estonian Committee on Bioethics and Human Research (Estonian Ministry of Social Affairs). All Estonian biobank participants have signed a broad informed consent form and the study was carried out under ethical approval 1.1-12/2856 from the Estonian Committee on Bioethics and Human Research (Estonian Ministry of Social Affairs) and released under data release P03.

Summaries of all posterior distributions obtained and the MLMA associated results are deposited on Dryad: Robinson, Matthew (2021), Maximizing GWAS discovery and genomic prediction accuracy in Biobank data, Dryad, Dataset, https://doi.org/10.5061/dryad.gtht76hmz

## Code availability

GMRM software is fully open source and available at: https://github.com/medical-genomics-group/gmrm.

## Supplementary Material

**Algorithm 1:**
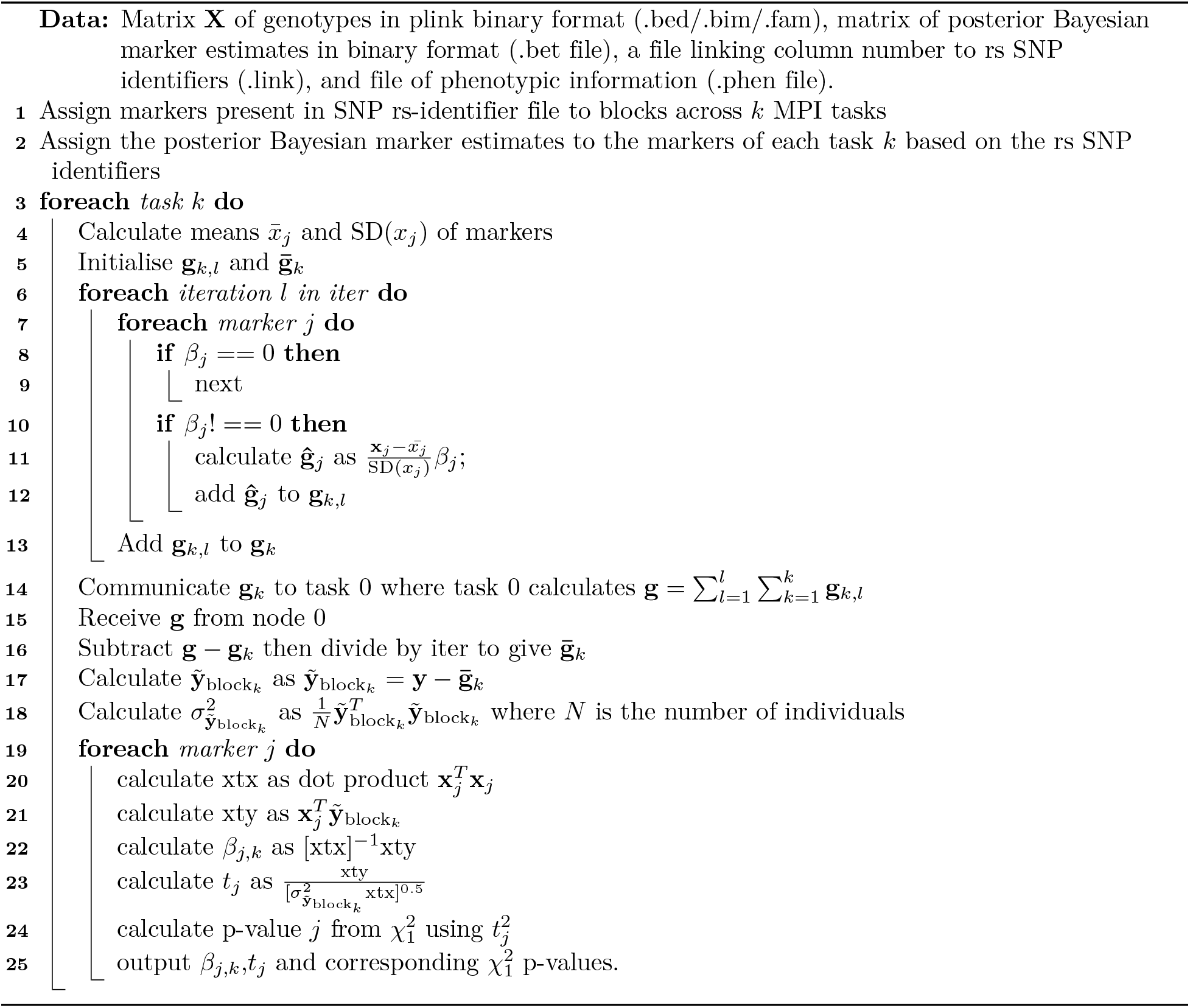
GMRM MLMA algorithm.

**Table S1.**
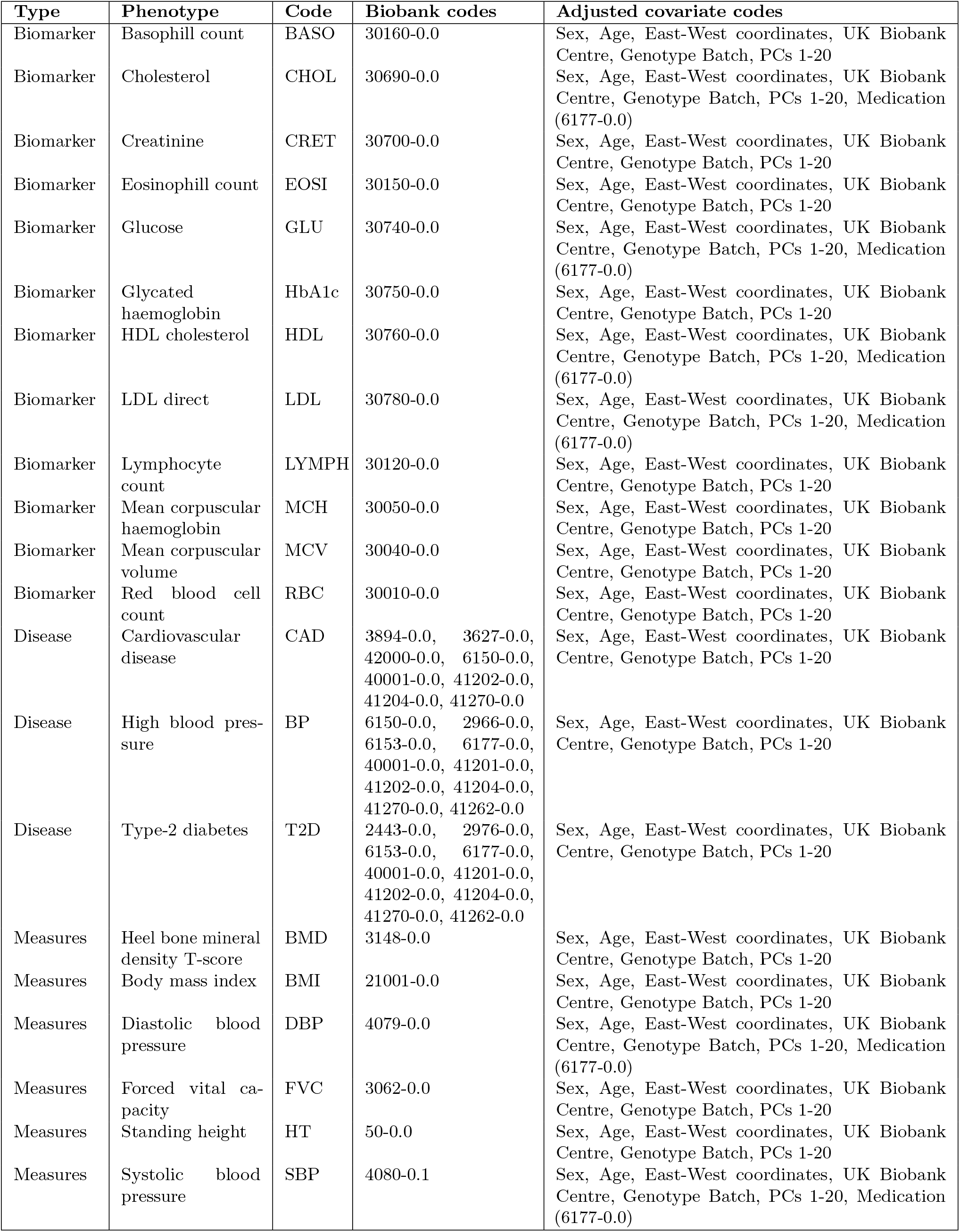
UK Biobank phenotypes used within the study. Columns of the table give the type of trait, the name, the trait code used for the figures, the UK Biobank codes used to construct the trait values, and the covariates adjusted for within the analysis.

| Type | Phenotype | Code | Biobank codes | Adjusted covariate codes |
| --- | --- | --- | --- | --- |
| Biomarker | Basophil count | BASO | 30160-0.0 | Sex, Age, East-West coordinates, UK Biobank Centre, Genotype Batch, PCs 1-20 |
| Biomarker | Cholesterol | CHOL | 30690-0.0 | Sex, Age, East-West coordinates, UK Biobank Centre, Genotype Batch, PCs 1-20, Medication (6177-0.0) |
| Biomarker | Creatinine | CRET | 30700-0.0 | Sex, Age, East-West coordinates, UK Biobank Centre, Genotype Batch, PCs 1-20 |
| Biomarker | Eosinophil count | EOSI | 30150-0.0 | Sex, Age, East-West coordinates, UK Biobank Centre, Genotype Batch, PCs 1-20 |
| Biomarker | Glucose | GLU | 30740-0.0 | Sex, Age, East-West coordinates, UK Biobank Centre, Genotype Batch, PCs 1-20, Medication (6177-0.0) |
| Biomarker | Glycated haemoglobin | HbA1c | 30750-0.0 | Sex, Age, East-West coordinates, UK Biobank Centre, Genotype Batch, PCs 1-20 |
| Biomarker | HDL cholesterol | HDL | 30760-0.0 | Sex, Age, East-West coordinates, UK Biobank Centre, Genotype Batch, PCs 1-20, Medication (6177-0.0) |
| Biomarker | LDL direct | LDL | 30780-0.0 | Sex, Age, East-West coordinates, UK Biobank Centre, Genotype Batch, PCs 1-20, Medication (6177-0.0) |
| Biomarker | Lymphocyte count | LYMPH | 30120-0.0 | Sex, Age, East-West coordinates, UK Biobank Centre, Genotype Batch, PCs 1-20 |
| Biomarker | Mean corpuscular haemoglobin | MCH | 30050-0.0 | Sex, Age, East-West coordinates, UK Biobank Centre, Genotype Batch, PCs 1-20 |
| Biomarker | Mean corpuscular volume | MCV | 30040-0.0 | Sex, Age, East-West coordinates, UK Biobank Centre, Genotype Batch, PCs 1-20 |
| Biomarker | Red blood cell count | RBC | 30010-0.0 | Sex, Age, East-West coordinates, UK Biobank Centre, Genotype Batch, PCs 1-20 |
| Disease | Cardiovascular disease | CAD | 3894-0.0, 3627-0.0, 42000-0.0, 6150-0.0, 40001-0.0, 41202-0.0, 41204-0.0, 41270-0.0 | Sex, Age, East-West coordinates, UK Biobank Centre, Genotype Batch, PCs 1-20 |
| Disease | High blood pressure | BP | 6150-0.0, 2966-0.0, 6153-0.0, 6177-0.0, 40001-0.0, 41201-0.0, 41202-0.0, 41204-0.0, 41270-0.0, 41262-0.0 | Sex, Age, East-West coordinates, UK Biobank Centre, Genotype Batch, PCs 1-20 |
| Disease | Type-2 diabetes | T2D | 2443-0.0, 2976-0.0, 6153-0.0, 6177-0.0, 40001-0.0, 41201-0.0, 41202-0.0, 41204-0.0, 41270-0.0, 41262-0.0 | Sex, Age, East-West coordinates, UK Biobank Centre, Genotype Batch, PCs 1-20 |
| Measures | Heel bone mineral density T-score | BMD | 3148-0.0 | Sex, Age, East-West coordinates, UK Biobank Centre, Genotype Batch, PCs 1-20 |
| Measures | Body mass index | BMI | 21001-0.0 | Sex, Age, East-West coordinates, UK Biobank Centre, Genotype Batch, PCs 1-20 |
| Measures | Diastolic blood pressure | DBP | 4079-0.0 | Sex, Age, East-West coordinates, UK Biobank Centre, Genotype Batch, PCs 1-20, Medication (6177-0.0) |
| Measures | Forced vital capacity | FVC | 3062-0.0 | Sex, Age, East-West coordinates, UK Biobank Centre, Genotype Batch, PCs 1-20 |
| Measures | Standing height | HT | 50-0.0 | Sex, Age, East-West coordinates, UK Biobank Centre, Genotype Batch, PCs 1-20 |
| Measures | Systolic blood pressure | SBP | 4080-0.1 | Sex, Age, East-West coordinates, UK Biobank Centre, Genotype Batch, PCs 1-20, Medication (6177-0.0) |

**Figure S1.**
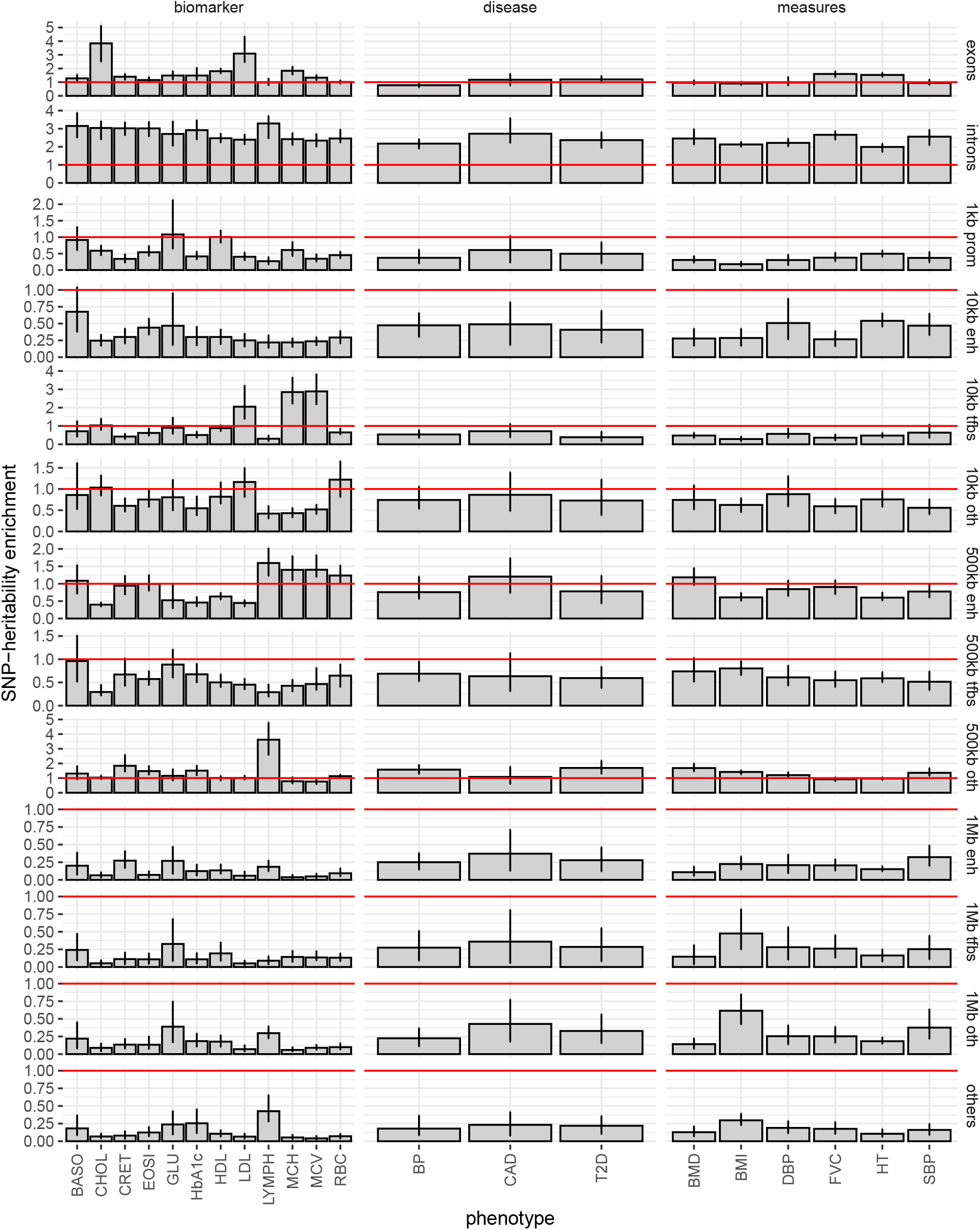
SNP-heritability enrichment estimated by GMRM. SNP heritability enrichment estimates calculated as the proportion of SNP-heritability attributable to each annotation group divided by the proportion of markers in the model given the total number of markers. If the average effect sizes of markers within a given annotation are larger than expected given the number of markers entering the model for that annotation then the value obtained should be greater than 1 (the red line shown). Conversely, smaller than expected marker effects will yield values less than 1.Error bars in give 95% credible intervals. Full trait codes are given in Supplementary Table 1. exons = SNPs located in exonic regions, introns = SNPs located in intronic regions, prom = SNPs located in promotors, tfbs = SNPs located in transcription factor binding sites, enh = enhancers, oth = markers not mapping to a functional group but to a location from a nearest gene, others = markers not within 1Mb of a gene and that have no known regulatory function.

**Figure S2.**
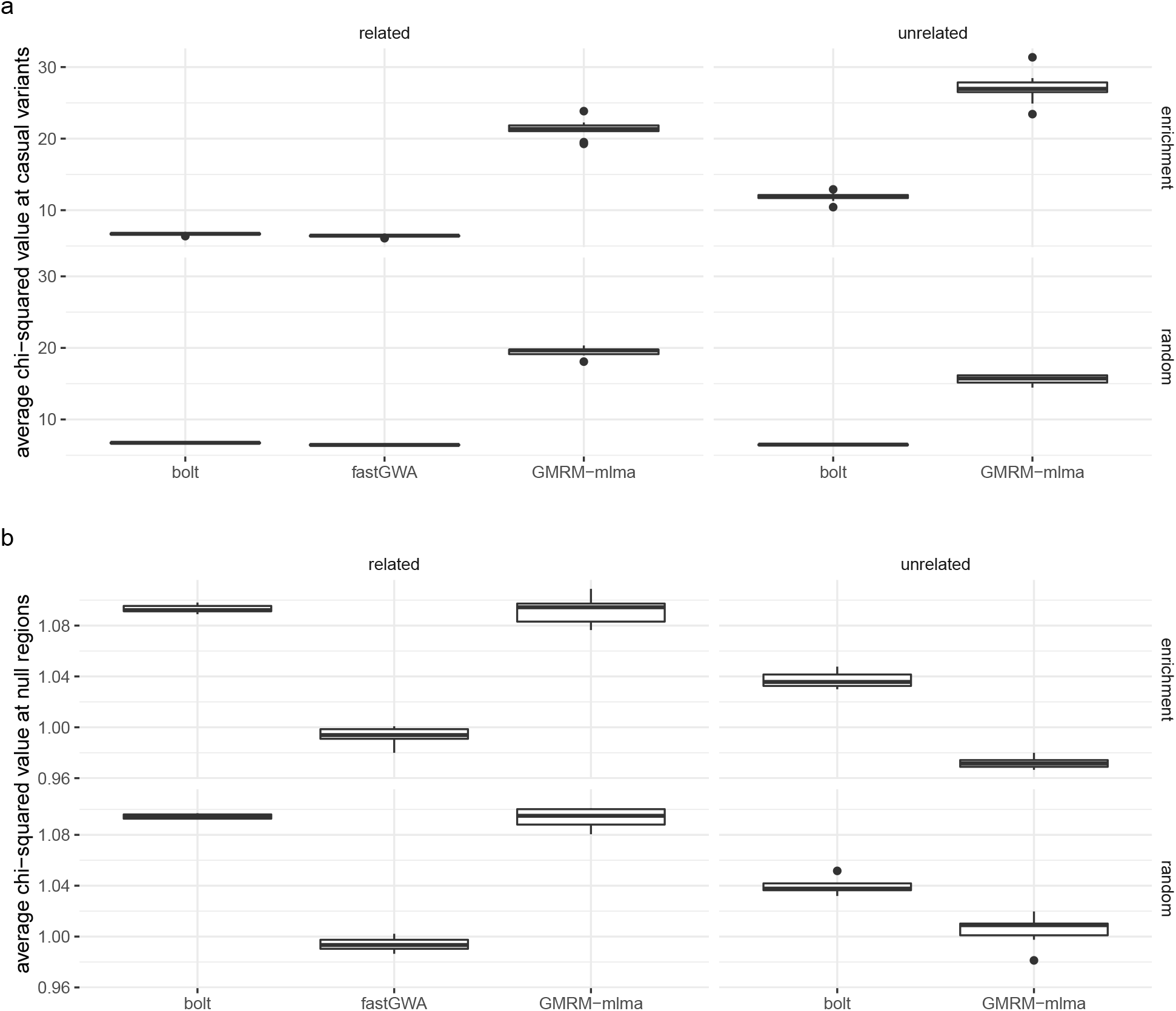
Simulation study for GMRM. (a) The y-axis gives the average *χ*^2^ values for the approximate mixed linear model test statistics at the simulated causal variants as compared to models run using fastGWA and bolt-LMM. The plot is faceted into two columns representing phenotypes that were simulated using either a mixture of 10,000 randomly sampled UK Biobank sibling pairs and 80,000 randomly sampled unrelated UK Biobank individuals (related), or 100,000 randomly sampled UK Biobank individuals with SNP marker relatedness estimates <0.2 (unrelated). The plot is also faceted into two rows representing 10,000 causal variants from chromosomes 1, 3, 5, 7 and 9 that were either randomly sampled with effect sizes drawn from a normal distribution with zero mean variance 0.5/10000 (random) or were sampled differently across genomic annotations creating effects size differences acorss genomic groups (see Methods). Boxplots show the distribution of values obtained for ten simulation replicates. (b) shows the same simulation settings but the average *χ*^2^ values for the approximate mixed linear model test statistics are given for the null chromosomes 2, 4, 6, 8, and 10 where no causal variants were simulated.

**Figure S3.**
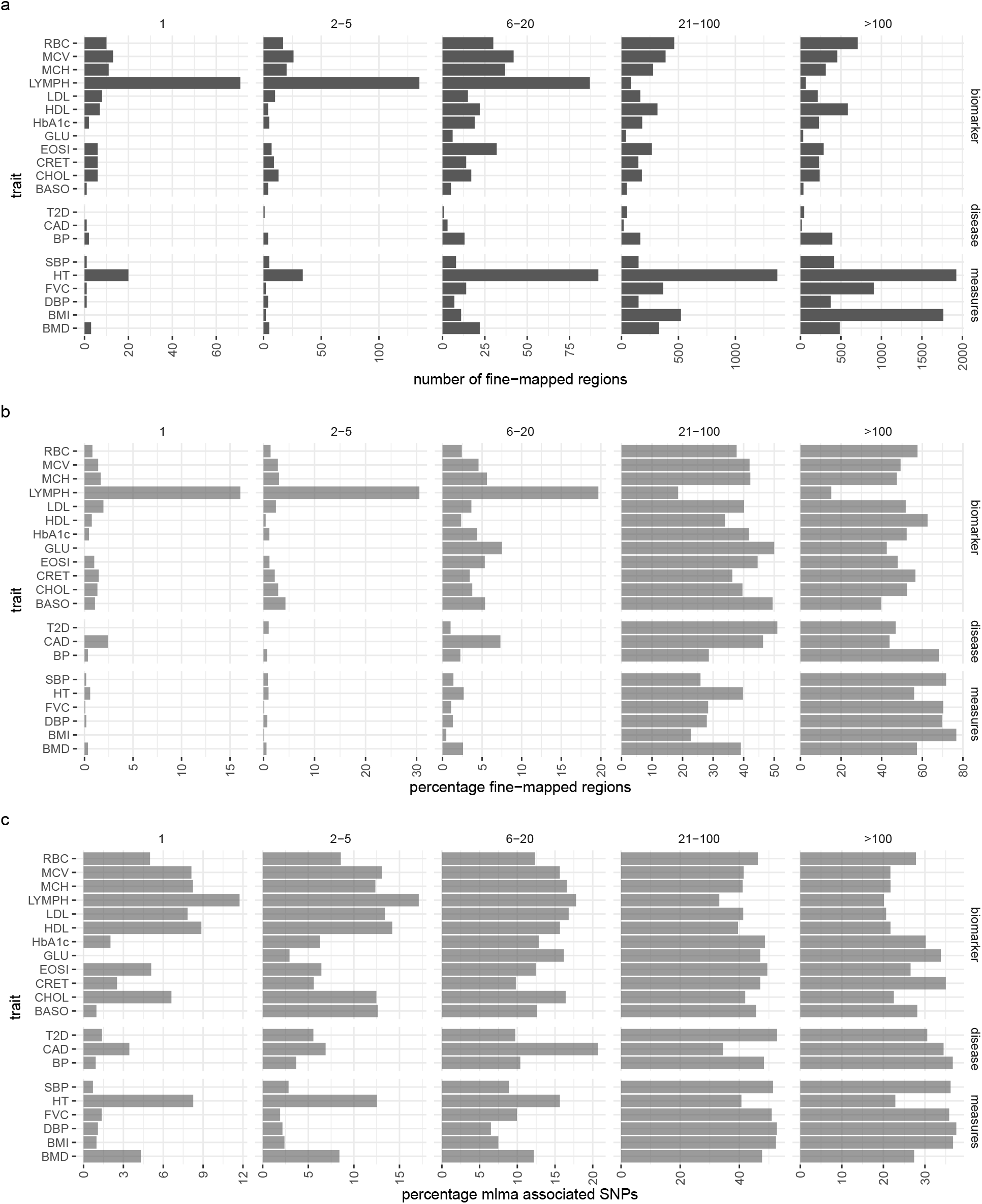
GMRM Bayesian fine-mapping approach to localise the inclusion of markers in the model into SNP sets based on their LD. (a) The trait acronyms are given on the y-axis and the plot is faceted into five columns representing set of SNPs that are correlated at LD *R*^2^ ≥ 0.1, from some genomic regions having only a single SNP in LD with no others, to ≥ 100 markers in LD. Barplots give a count of the number of SNP sets for which the posterior probability of explaining greater than 0.001% of the phenotypic variance was greater than 95%. (b) For each trait, the percentage of SNP sets for which the posterior probability of explaining greater than 0.001% of the phenotypic variance was greater than 95% for different sizes of SNP set. (c) GMRM-MLMA significantly associated SNPs grouped into the same SNP sets based on LD. Across traits, a lower percentage of associations fine-map to either single SNPs, or to groups of five or less SNPs in LD as compared to the percentage of the discoveries from the GMRM-MLMA approach.

**Figure S4.**
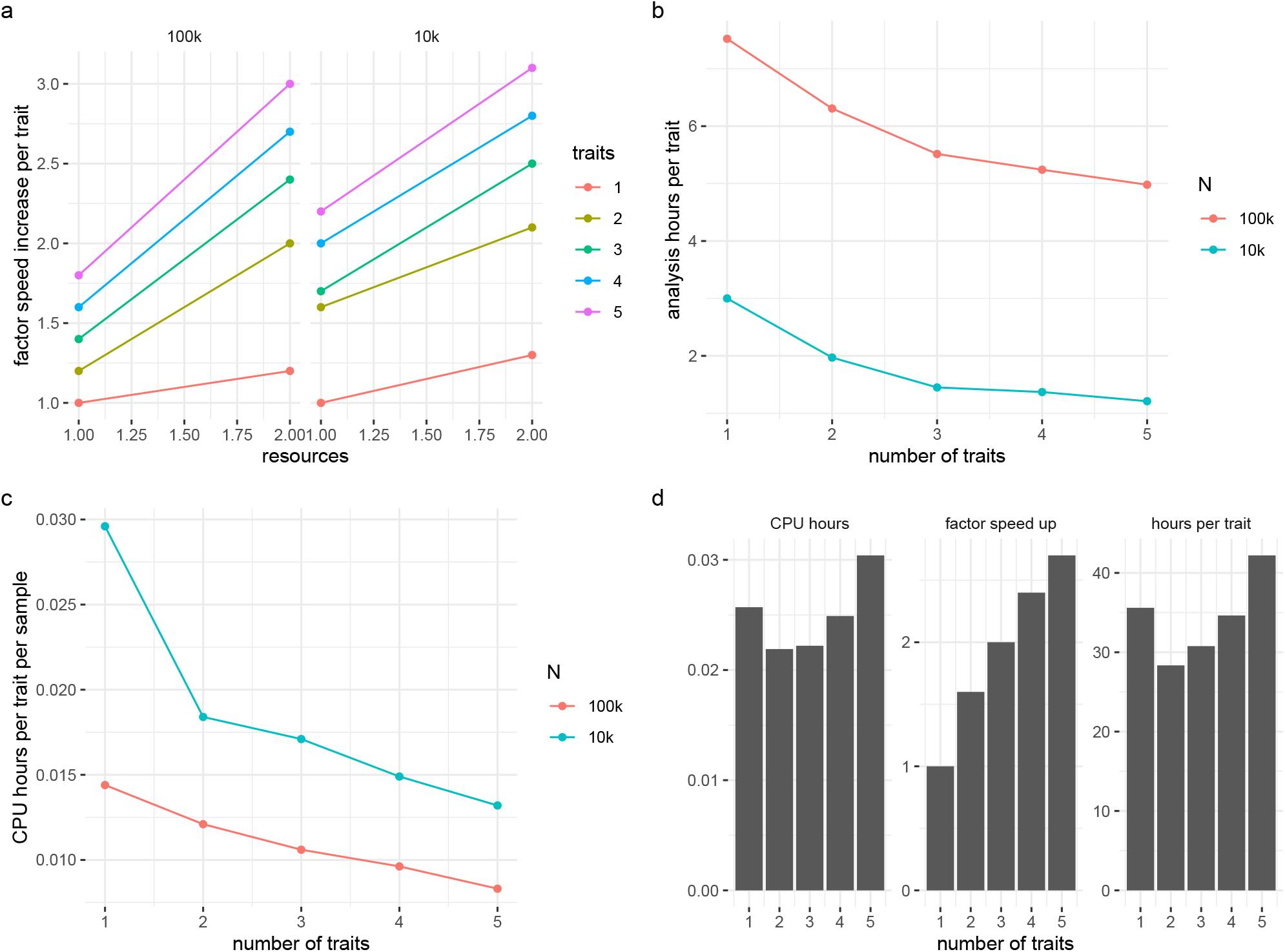
Scaling of GMRM software for an analysis of 2.2M markers. (a) For two resource settings of (1) 12 MPI processes each using 4 CPU cores on a single computer and 24 MPI processes each using 4 CPU cores across two computers, labelled as 1 and 2 on the x-axis, the factor speed-up per trait analysed is given on the y-axis. Speed-up factors are presented relative to the time taken to estimate 5,000 draws from the posterior distribution of a single trait in resource setting 1 (baseline time). We calculated the speed-up times for data from 10,000 individuals (10k) and 100,000 individuals (100k) separately. For example, for the analysis of 5 traits at the same time for 10k setting in resource setting 2 the value presented was calculated as: *t*_1,1_*/*(*t*_5,2_*/nt*), where *t*_1,1_ is the time at baseline for 1 trait in resource environment 1, *t*_5,1_ is the time for analysing 5 traits simultaneously in resource environment 2, and *nt* is the number of traits analysed. Thus we obtain the proportional increase in time obtained relative to the baseline. (b) The number of hours required to obtain 5,000 posterior samples at two different sample sizes, decreases steeply but then begins to tail-off with a large number of simultaneously analysed traits. (c) In the configurations presented here, our algorithm displays sub-linear scaling of CPU core hours per trait per sample, with improved CPU use efficiency as sample size and number of traits analysed increases, essentially becoming more efficient as sample size and the number of traits grows. These examples were specifically designed to remain within memory and last level memory cache restrictions of the high-performance compute hardware used here in order to demonstrate the performance of our algorithm. Mostly, with the multi-trait version some MPI communication is shared among the traits, explaining the sub-linear scaling observed here. (d) In our hardware, at 400,000 individuals, 2.2M markers, and 72 MPI processes last-level cache size overflowed at greater than two traits analysed simultaneously, resulting in increased CPU hours per trait per sample (CPU hours), reduced speed-up, and an increase in the analysis time per trait. Thus, performance gains from our algorithm will become hardware limited. If the epsilon vectors stay in the cache we get continually improved performance as shown in (a) through (c). However, with more individuals, more traits, and more tasks per socket the hardware limits the sub-linear scaling to the point where degraded performance is observed as compared to single trait processing. The main considerations are the last-level cache size, the number of individuals and the number of traits to be analysed, and we advise users to set 1 MPI task per socket and estimate how many epsilon vectors can be held in the last level cache of the CPU.

**Figure S5.**
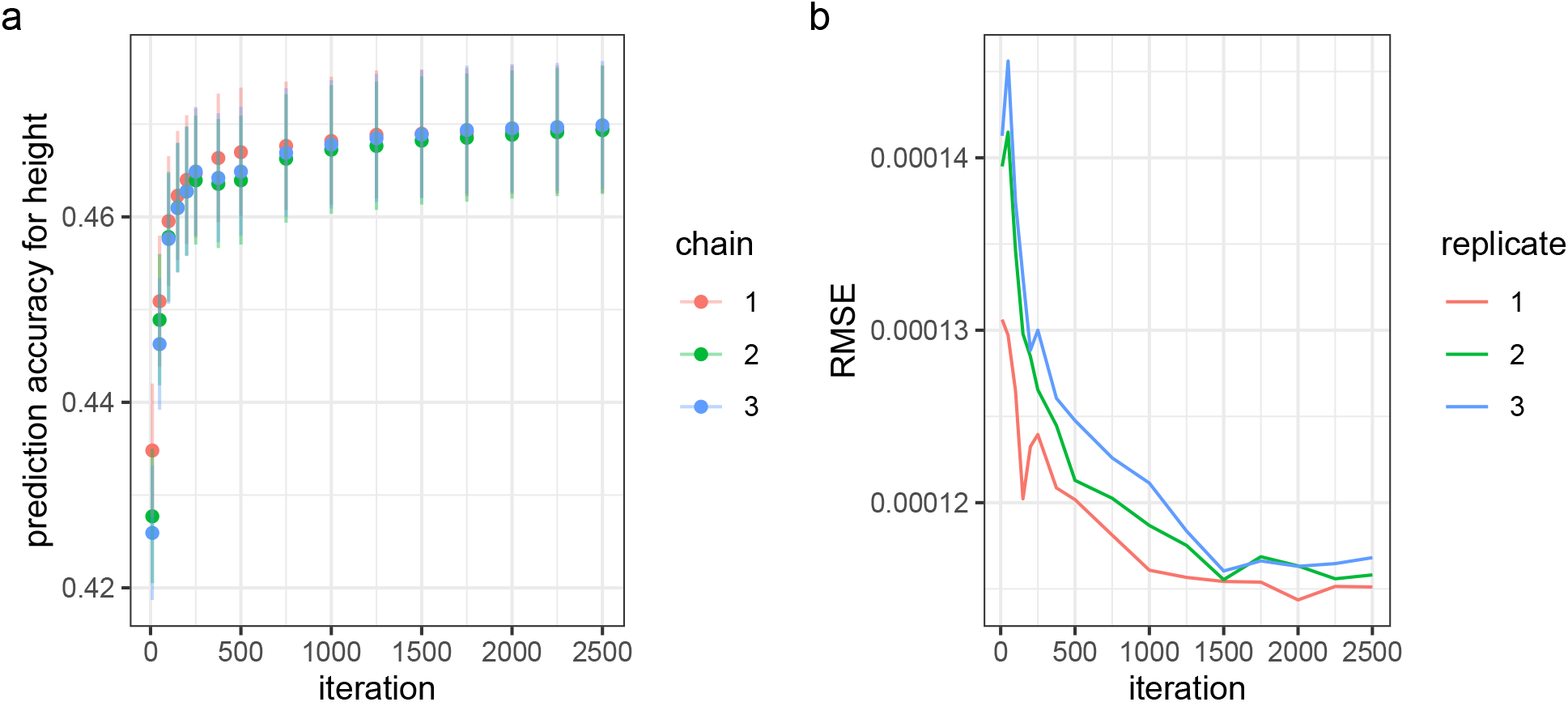
Convergence of GMRM software. (a) Convergence of prediction accuracy in a hold out sample of 30,000 UK Biobank individuals for human height across three independent bulk synchronous Gibbs chains with synchronisation occurring after the update of 20 markers across 60 MPI process for 2.17 million SNP markers and 428,747 individuals. (b) The root mean square error of the marker effect size estimates of randomly assigned LD independent causal variants in simulation study of 100,000 unrelated individuals from the UK Biobank for three replicates.

